# Novel insights into the genome organization of *Rhizobiaceae*: identification of linear plasmids

**DOI:** 10.1101/2025.05.29.656789

**Authors:** Nemanja Kuzmanović, Ryan R. Wick, Michał Kalita, Joanna Puławska, George C. diCenzo, Michael F. Hynes, Kornelia Smalla, Doreen Babin, Torsten Thünen

## Abstract

2.

Members of the family *Rhizobiaceae* typically have multipartite genomes, that are split between two or more replicons, including the chromosome and a variable number of extrachromosomal replicons (chromids and plasmids). Nearly all *Rhizobiaceae* replicons sequenced and described to date have a circular topology, with the exception of the linear chromid found in the genomes of most *Agrobacterium* spp. In this study, genomes of five nonpathogenic *Agrobacterium* strains and one plant tumorigenic *Allorhizobium* strain were fully sequenced. Surprisingly, genome analysis revealed that these six strains each carry an 80-kbp linear plasmid. Linear plasmids were so far not identified in this bacterial family or other bacteria within the class *Alphaproteobacteria*. The ends of all six plasmids identified in this study have a hairpin structure with covalently closed ends. The plasmid sequences showed a high degree of homology, clearly indicating their common ancestry. Database searches led to the identification of additional linear plasmids in previously published *Rhizobiaceae* genome assemblies that were not previously recognized to have linear plasmids, suggesting that these replicons may be more widespread. Most likely, linear plasmids may be even more widely distributed than anticipated. Although the biological functions of the linear plasmids identified in this study remain unknown, they are associated with both nonpathogenic and plant tumorigenic *Rhizobiaceae* strains.

**Impact statement:** The family *Rhizobiaceae* includes some remarkable and important representatives, such as plant symbiotic bacteria (rhizobia) and plant pathogenic bacteria associated with neoplasia (agrobacteria). In this study, the complete genome sequences of six *Rhizobiaceae* strains were generated and their genome organizations were examined. Strikingly, our results showed that these six strains harbor a linear plasmid. Moreover, GenBank searches suggested that linear plasmids may be even more widespread in the family *Rhizobiaceae*. Linear plasmids may go undetected in genome sequencing studies if the assemblies are not specifically examined for linear plasmid. Overall, this study provides further evidence for the extraordinary genome plasticity of members of the family *Rhizobiaceae* and expands the taxonomic range in which linear plasmids have been identified. To the best of our knowledge, this is the first report of linear plasmids in the family *Rhizobiaceae* or the class *Alphaproteobacteria*.

**Data summary:** The whole-genome sequences have been deposited at DDBJ/ENA/GenBank under the accessions CP192696-CP192701 (Av2), CP000000-CP000000 (rho-7.1), CP000000-CP000000 (rho-8.1), CP000000-CP000000 (rho-11.1), CP000000-CP000000 (rho-13.3), and CP000000-CP000000 (rho-14.1), within the BioProjects PRJNA557463 and PRJNA1009994. The raw sequencing reads were deposited in the Sequence Read Archive (SRA) under the same BioProjects PRJNA557463 and PRJNA1009994: https://www.ncbi.nlm.nih.gov/bioproject/PRJNA557463 and https://www.ncbi.nlm.nih.gov/bioproject/PRJNA1009994. NCBI submission for five genome sequences is undergoing processing and accession numbers will be added when available; in https://figshare.com/s/29a9e621adc1b66d0957

## 5. Introduction

The *Rhizobiaceae* is a large family of Gram-negative bacteria, classified within the alphaproteobacterial order *Hyphomicrobiales* (*Rhizobiales*) [1]. This family hosts diverse bacteria occurring in various environments, including plants, soil, sediments, and water [2]. The most remarkable and well-known *Rhizobiaceae* are plant symbiotic bacteria (rhizobia) that primarily belong to the genera *Rhizobium*, *Sinorhizobium*, *Allorhizobium*, *Pararhizobium*, *Neorhizobium* and *Shinella* [3], as well as plant pathogenic bacteria (agrobacteria) that are predominantly found within the genera *Agrobacterium*, *Allorhizobium* and *Rhizobium* [4–6].

Typically, the genomes of members of the family *Rhizobiaceae* are multipartite [7], meaning that they are split between two or more large (>350 kbp) DNA fragments or replicons [8]. In other words, their genome consists of the chromosome (primary replicon) and a variable number of extrachromosomal (secondary) replicons. Taxa of the family *Rhizobiaceae* are not an exception in the bacterial world, as roughly 10% of currently fully sequenced bacterial genomes are multipartite [8]. In multipartite genomes, the chromosome is the largest replicon that carries most of the genes involved in core cellular processes, which are termed essential or core genes. Extrachromosomal replicons can be variable in their size and indispensability, leading to the emergence of classification schemes to distinguish them [8]. One such scheme classifies extrachromosomal replicons into the following five groups: second chromosome, chromid, megaplasmid and plasmid [8]. In this respect, a second chromosome is defined as a replicon that result from the split of the chromosome into two replicons. A chromid is a replicon that carries essential (or at least very important for survival) genes and has evolved from plasmids, and therefore has both plasmid and chromosome characteristics [9]. Megaplasmids and plasmids carry no core genes and are therefore nonessential replicons, although they are vitally important for the survival of bacteria in some environments. They are distinguished based on their size, although the boundaries between megaplasmids and plasmids are arbitrary. For instance, 8 [8] proposed a cutoff of 350 kb, where the replicons larger than 350 kb are classified as megaplasmids, while others have pointed out that a strict size threshold across taxa is likely not appropriate [10].

Moreover, bacterial replicons can be also distinguished based on their topology (circularity or linearity). In the family *Rhizobiaceae*, nearly all replicons sequenced and described to date have a circular topology. The exception is in the genus *Agrobacterium*, whose most representatives carry a linear chromid, in addition to the primary circular chromosome and a variable number of circular plasmids [11–13]. The linear chromids of the genus *Agrobacterium* have covalently closed hairpin ends, which are generated by the activity of a protelomerase (telomere resolvase) [14]. The only representative of this genus that carry no linear chromid is a recently described species *Agrobacterium divergens*, belonging to a remote *Agrobacterium* clade [15].

In this study, the complete genome sequences of five *Agrobacterium* strains and one *Allorhizobium* strain were generated and their genome organizations were assessed. Unlike other members of the family *Rhizobiaceae*, our results indicated that these six strains carry a linear plasmid. Linear plasmids are generally associated with a limited number of bacteria, including some members of the class *Actinomycetes* (e.g., *Streptomyces* and *Rhodococcus*) [16], *Borrelia* [17], and *Enterobacteriaceae* (e.g., *Salmonella* and *Klebsiella*) [18,19]. To the best of our knowledge, this is the first report of linear plasmids in the family *Rhizobiaceae*, or even more broadly, within the order *Alphaproteobacteria*.

## 6. Methods

### 6.1 Bacterial strains

Six *Rhizobiaceae* strains were analyzed in this study (Table 1). *Allorhizobium* sp. Av2 was isolated from a crown gall tumor on grapevine and carries a tumor-inducing (Ti) plasmid (as determined by PCR) responsible for pathogenicity on plants [20]. The five *Agrobacterium* strains were isolated from aerial crown gall tumors on rhododendron [21]. They do not carry a Ti plasmid and are nonpathogenic. However, as previously determined by analysis of their draft genomes, they carry an opine-catabolic plasmid enabling them to thrive in crown gall tumors [21]. Strains Av2, rho-8.1 and rho-13.3 belong to novel, as-yet undescribed species [6,21] and are assigned here to the genus level (Table 1). All bacterial cultures were preserved at -80°C in nutrient broth with 20% glycerol.

**Table 1.**
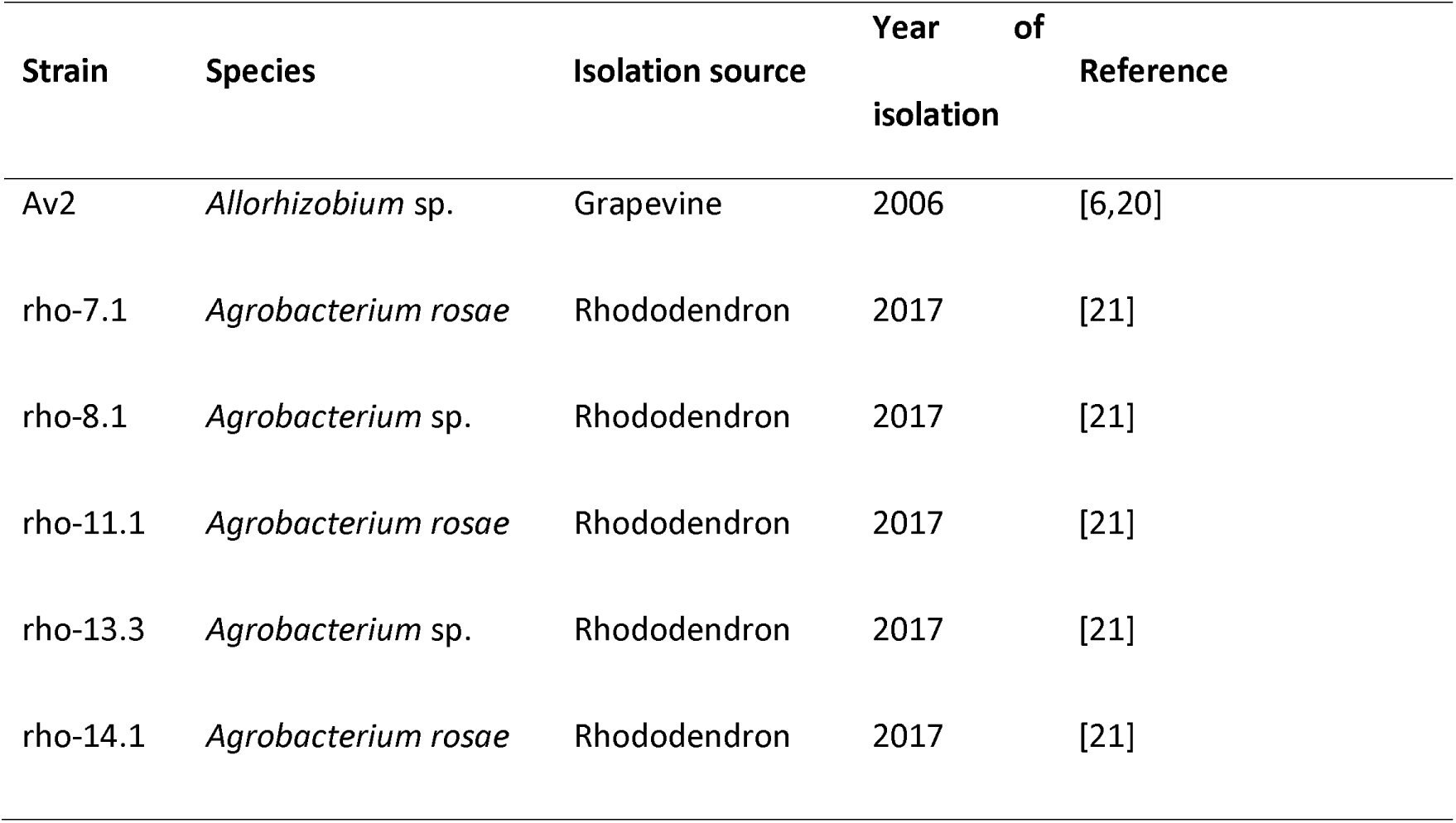
*Rhizobiaceae* strains analyzed in this study. All strains were isolated from crown gall tumors on plants.

### 6.2 Genomic DNA extraction

High molecular weight (HMW) genomic DNA (gDNA) was extracted from bacteria grown in nutrient broth (Difco BD, MD, USA), on nutrient agar, or on TY medium (tryptone 5 g/L, yeast extract 3 g/L, CaCl_2_×2H_2_O 0.9 g/L, agar 16 g/L) at 28°C for 24-48 h using a Qiagen Genomic DNA Buffer Set (Qiagen, Germany; Cat. No. 19060) and Qiagen genomic tip 100/G gravity-flow, anion exchange columns (Cat. No. 10243). The integrity of the HMW gDNA was roughly assessed by electrophoresis in 0.8% agarose gel. The purity and approximate concentration of HMW gDNA was determined using a NanoDrop instrument. Additionally, the quantity of HMW gDNA was checked using a Qubit dsDNA Broad Range (BR) Assay Kit (Thermo Fisher Scientific, Waltham, MA, USA).

### 6.3 Nanopore sequencing and initial data processing

For strain rho-13.3, the DNA library was constructed using a ligation sequencing kit (SQK- LSK110; Oxford Nanopore Technologies [ONT]), with no size selection, and sequenced on a MinION Mk1B with a R9.4.1 flow cell (FLO-MIN106D). The raw FAST5 files were basecalled using Guppy v6.4.6 (ONT) with a super accurate (SUP) model (dna_r9.4.1_450bps_sup.cfg) and the “--trim_adapters” option. The resulting FASTQ files were filtered using Filtlong v0.2.1 (https://github.com/rrwick/Filtlong) with the options --min_length 6000 and -- target_bases 1000000000 or 2000000000.

For the remaining strains, DNA libraries were prepared using ONT Native Barcoding Kit 24 V14 (SQK-NBD114.24), with no size selection, and sequenced on a MinION Mk1B with a R10.4.1 flow cell (FLO-MIN114). The raw POD5 files were basecalled using Dorado v0.7.1 (ONT) with a super accurate (SUP) model (dna_r10.4.1_e8.2_400bps_sup@v5.0.0). Barcode demultiplexing was also performed using Dorado. The resulting BAM files were converted to FASTQ format using SAMtools v1.18 [22]. The FASTQ files were filtered using Filtlong (with options --min_length 6000 and --keep_percent 90, in two subsequent runs), as suggested by 23 [23].

### 6.4 Genome assembly, error-correction and annotation

Genome assembly was performed with a hybrid approach, using both long reads generated in this study and short reads generated in our previous studies [6,21]. Primarily, we relied on the long-read-first hybrid assembly, which involved long-read assembly, followed by long-read polishing and short-read polishing. In particular, long-read assembly was primarily done using Flye v2.9.4-b1799 [24]. In some cases, additional assemblers were used, including Raven v1.8.3 [25], Miniasm/Minipolish v0.3-r179/v0.1.2 [26–28], Canu v2.2 (with option “genomeSize=6.5m”) [29], Trycycler v0.5.5 [30] and/or Autocycler v0.1.0 [31]. The resulting assemblies were manually curated and processed to assess if the entire genome and all the replicons were represented. The presence of plasmids was additionally assessed by gel electrophoresis methods (see below). Long-read polishing of the resulting raw assemblies was achieved using Medaka 1.11.3 (medaka_consensus; ONT), while the subsequent short-read polishing was conducted using Polypolish v0.6.0 [32]. Automated long-read first hybrid assembly was also performed by Hybracter v0.7.3 [33], using raw non-filtered reads as an input and option “-c 3000000”. Moreover, for comparison and for eventual identification of small plasmids, short-read first hybrid assembly was carried out using Unicycler v0.5.1 and Spades v4.0.0 [34]. All assemblers were run with standard parameters if not otherwise stated. Finally, to correct any remaining errors, long and short reads were mapped to assemblies with minimap2 v2.28-r1209 [35] and BWA v0.7.18-r1243- dirty [36], respectively, and the resulting read alignments were visually inspected. For the examination of assembly graphs produced by some assemblers, the software Bandage v0.8.1 [37] was used.

The final assemblies were annotated using the NCBI Prokaryotic Genomes Annotation Pipeline (PGAP) v2024-07-18.build7555 [38]. Additionally, for annotation of the linear plasmids, eggNOG-mapper version emapper-2.1.12 [39] and the eggNOG orthology data [40] were used. For eggNOG-mapper, sequence searches were performed using DIAMOND version 2.0.11 [41]. Moreover, to aid functional annotation of some loci, BLASTp comparisons against the NCBI non-redundant (nr) protein database (https://blast.ncbi.nlm.nih.gov/Blast.cgi; last accessed on April, 2025) [42] were conducted.

### 5.5 Sequence and phylogenetic analysis

Synteny plots were made using the pgv-mummer workflow, which is a part of the pyGenomeViz version 1.5.0 genome visualization python package (https://github.com/moshi4/pyGenomeViz). This workflow employs MUMmer version 3.23 [43] to align sequences. The sequences were aligned at the nucleotide level.

Whole-plasmid average amino acid identity (AAI) was computed using EzAAI version 1.2.3 [44], including the dependencies Prodigal version 2.6.3 [45] and mmseqs2 version 16.747c6 [46]. The minimum identity threshold for AAI calculations was set to 0.2 (20%).

For phylogenetic analysis, a protelomerase protein multi-sequence alignment was generated using MAFFT version 7 [47]. A maximum likelihood (ML) phylogeny was inferred using IQ-TREE 1.6.12 [48] available through the IQ-TREE web server (http://iqtree.cibiv.univie.ac.at/) [49]. Model selection was conducted using IQ-TREE ModelFinder [50] based on Bayesian Information Criterion (BIC) [51]. Branch supports were assessed by ultrafast bootstrap analysis (UFBoot) [52] and the SH-aLRT test [53] using 1000 replicates. The trees were visualized using FigTree version 1.4.4 (https://github.com/rambaut/figtree) and edited using Inkscape version 1.2.1 (https://inkscape.org/).

### 6.6 PCR analysis

For experimental detection of the presence of the linear plasmids identified in the strains sequenced in this study, a specific primer set, consisting of linp-topo_f (5’- GCGTACTTGATCGGCTTGTT-3’) and linp-topo_r (5’- CGCCAACCTTTCGACTTCAA-3’), targeting a gene putatively encoding DNA topoisomerase I was designed using Primer 3.0 version 4.1.0 [54]. The size of the expected amplification product is 787 bp. The PCR amplifications were performed in 25 μL reaction mixtures containing 1× OneTaq Quick-Load Buffer (New England Biolabs), 200 μM of each dNTP, 0.2 μM of each primer, 0.5 U of OneTaq Quick-Load DNA Polymerase (New England Biolabs), and 10-20 ng template DNA. The thermal profile was as follows: initial denaturation at 94°C for 1 min; 35 cycles of denaturation at 94°C for 30 s, annealing at 55°C for 1 min, elongation at 68°C for 1 min; and a final extension at 68°C for 5 min. Amplification of the 16S rRNA gene was performed for DNA samples purified from the Eckhardt-type gels (see the next paragraph), as described before [55].

### 6.7 Eckhardt-type gel electrophoresis

Separation and visualization of bacterial replicons, primarily plasmids, was conducted using the modified method of Eckhardt [56]. For this purpose, we used two protocols (I and II). Protocol I was as described in our previous publication [4]. Protocol II was similar to one reported by Hynes et al. [57]. Electrophoresis was performed in a gel containing 0.7% (w/v) agarose and 0.3% (w/v) sodium dodecyl sulfate (SDS) in 1× Tris-borate-EDTA (TBE) buffer. SDS was added to the gel after melting the agarose as a 10% solution prepared in 1× TBE buffer. Bacteria were grown in 15 mL tubes in HP medium (4 g/L peptone, 0.5 g/L yeast extract, 0.5 g/L tryptone, 0.2 g/L CaCl_2_, 0.2 g/L Mg_2_SO_4_) on a rotary shaker (200 rpm) at 28°C for approximately 24 h. A 0.1-0.2 mL aliquot of bacterial culture at an optical density at 600 nm (OD_600_) of 0.3-0.5 was transferred to the microfuge tube and kept on ice. Half a mL of cold 0.3% (m/v) sodium lauroylsarcosinate solution (prepared in 1× TBE buffer) was added, after which the cell suspension was gently mixed by pipetting and then centrifuged at 10,000 g for 1 min. The bacterial pellet was resuspended in 20 µL of ice-cold lysis solution containing 10% [w/v] sucrose, 1 mg/mL lysozyme, and 0.4 mg/mL RNase A, freshly prepared in 1× TBE buffer. The mixture was then immediately loaded into wells of the gel. Electrophoresis was run first at room temperature at 0.4 V/cm for 30 min and then in the fridge or cold room (∼4°C) at 3.2 V/cm for 6-10 h. The gel was stained in an ethidium bromide solution (1 μg/mL) and the replicons were visualized under UV light. As markers, “*Agrobacterium fabrum*” C58^T^, *Allorhizobium ampelinum* S4^T^, and/or *R. rhizogenes* K84 carrying replicons of known size were used. For some gels, a Quick-Load 1 kbp Extend DNA Ladder (New England BioLabs, Inc., Ipswich, MA, USA) was also included.

### 6.8 Extraction of replicon DNA separated by Eckhardt-type gel electrophoresis

DNA was recovered from the respective replicons of interest visible on the gel after their separation by the Eckhardt-type method. Briefly, bands were cut out from the Eckhardt- type gels and DNA was purified using Zymoclean Large Fragment DNA Recovery kit (Zymo Research, CA, USA, Cat. No. D4045) following the manufacturer’s instructions. The only modification was that the amount of agarose dissolving buffer (ADB) was increased to 4-5 volumes per volume of the excised agarose gel slice. This modification was made to reduce the final concentration of SDS in the solution. SDS was present at a concentration of 0.3% (w/v) in the Eckhard-type agarose gel, while the indicated kit tolerance to this compound was ≤0.1%. To obtain a sufficient concentration of the replicon DNA, DNA from several lanes was cut out and combined before purification. The integrity of the recovered DNA was roughly assessed by electrophoresis using a 0.8% agarose gel. The purity and approximate concentration of DNA was determined using a NanoDrop instrument.

### 6.9 Illumina sequencing of replicon DNA, data processing and analysis

Library preparation and sequencing were performed by Novogene GmbH (Munich, Germany). Briefly, DNA libraries were prepared with the Novogene NGS DNA Library Prep Set (Cat. No. PT004), involving random DNA shearing into shorter fragments, end repair, A-tailing, and ligation with Illumina adapters. The resulting fragments with adapters were size selected, PCR amplified, and purified, after which they were sequenced using Illumina 150- bp paired-end sequencing on the Illumina NovaSeqX Plus platform (Illumina, CA, USA).

Adapter trimming and quality filtering of raw reads were conducted with fastp v0.23.4 [58] using default settings. The depth of coverage statistics (mean, max, and min) for each replicon was assessed as follows. First, “clean” reads were mapped to the corresponding complete genome sequence obtained in this study using BWA and the resulting BAM files were sorted with SAMtools “sort” command. Next, the read depth at each nucleotide position of the genome was calculated with the SAMtools “depth” option. Finally, mean, max, and min values for depth of coverage were calculated for each replicon using the datamash program (https://www.gnu.org/software/datamash/).

### 6.10 Pulsed-field gel electrophoresis

For preparation of agarose plugs, bacteria were grown in YEM medium (0.5 g/L yeast extract, 5 g/L mannitol, 0.5 g/L K_2_HPO_4_, 0.2 g/L Mg_2_SO_4_, 0.1 g/L NaCl) for 48 h at 28°C. A 5 ml culture was centrifuged (13 000 g for 10 min), after which the pellet was washed with 500 µL of 0.5 M NaCl, centrifuged again (13 000 g for 10 min), the supernatant removed, and the pellet resuspended in 500 µL of TE buffer. A 2% (w/v) low gelling temperature agarose (Sigma-Aldrich) was dissolved in TE buffer and kept in a water bath at 45°C for about 10 minutes to stabilize the temperature. The 500 µL bacterial suspension in TE buffer was mixed with 500 µL of melted low gelling temperature agarose, mixed thoroughly, and applied to wells (one 10-well plug mold = 10 plugs of the same bacterial strain). The mixture was left at room temperature for 10 minutes and then incubated at 4°C for a further 10 min to solidify. Then, the agarose plugs were pushed out from the wells into small plastic containers with closure. Ten ml of TE buffer with lysozyme (Merck) (1.5 mg lysozyme/ml TE buffer) was added to each container, and the plugs were incubated overnight at 37°C with gentle shaking (80 rpm).

The next day, the buffer was removed, and 10 ml of lysis buffer [50 mM EDTA (pH 8.0), 50 mM Tris-HCl (pH 8.0), 1% (w/v) N-lauryl sarcosine] with proteinase K (Sigma-Aldrich) (0.5 mg proteinase/ml buffer) was added. The plugs were incubated at 37°C with gentle shaking (80 rpm) for 48 hours.

Afterwards, the buffer was removed, 10 ml of TE buffer was added, and the plugs were shaken (60 rpm) at 37°C for 20-30 min. The buffer was removed, replaced with fresh TE buffer, and the incubation was repeated. To inactivate proteinase K, the TE buffer was removed, replaced with TE buffer containing phenylmethanesulfonyl fluoride (PMSF, Sigma-Aldrich) (0.4 mg PMSF/ml TE buffer), and incubated on a shaker with gentle mixing at 37°C for 1 hour. Then, the buffer was removed, and the plugs were washed twice with TE buffer – 10 ml at 37°C for 20 minutes each, with gentle mixing. Finally, the plugs were stored in TE buffer at 4°C until further analysis.

Gels for electrophoresis were prepared with 1% agarose in 0.5x TBE buffer. The agarose plugs were placed in wells and immobilized by overlaying with melted low gelling temperature agarose. Pulsed field gel electrophoresis (PFGE) was carried out in 0.5x TBE buffer using a CHEF-DR III system (Bio-Rad) and the following conditions: voltage 6 V/cm, reorientation angle 120°, switch time 1-25 s, 5-30 s or 10-40 s, temperature 14°C, time 22 h or 30 h.

For S1 nuclease treatment, the agar plugs were pre-incubated in 200 µL of 1x S1 buffer at room temperature for 30 min. Then, the plugs were incubated in 200 µL of fresh 1x S1 buffer containing 10 U of S1 nuclease (Thermo Fisher Scientific), and incubated at 37°C for 20 min. The reaction was stopped by suspending the plugs in 200 µL of 0.5M EDTA. The plugs in EDTA were incubated at room temperature for 10 min. Then, the EDTA solution was substituted with 1x TE and left at room temperature for at least 30 min prior to loading the gel.

As markers, *All. ampelinum* S4^T^ carrying replicons of known size and/or ProMega-Markers® Lambda Ladders (Promega) were also included.

## 7. Results and discussion

### Whole genome sequencing revealed the presence of a linear plasmid

A draft genome sequence of *Agrobacterium* sp. rho-13.3 was obtained in our previous study, and its phylogenetic position was elucidated [21]. As this strain formed a distinct and novel phylogenetic lineage of the genus *Agrobacterium*, which represented an outgroup to the other members of the *Agrobacterium* clade “rubi”, here we used ONT long-read sequencing to generate a higher quality genome assembly.

As the strain rho-13.3 was sequenced alone on an ONT flow cell, a tremendous amount of ONT data (9.4 Gbp) was generated, resulting in an excessively high depth of genome coverage (1640×; Table S1). Therefore, in this case, the data were filtered to keep approximately 1 or 2 Gbp of best reads, while still maintaining a high depth of genome coverage (∼175 and ∼350×) and a read length N50 of 45 or 41 kbp, respectively (Table S1).

The genome assembly of the strain rho-13.3 comprised seven replicons (Table 2), including one large circular chromosome and a chromid with a linear topology, which is typical for this bacterial genus. Additionally, it carried multiple circular plasmids, including an opine-catabolic (OC) plasmid as characterized previously [21]. However, the assembly of two circular plasmids proved difficult for strain rho-13.3. Whole-genome assembly was initially performed using the Flye assembler with the option “--nano-hq”. However, assemblies differed depending on which input reads were used. For instance, when the best 1 Gbp of reads (>6 kbp read length) were used as an input, two small circular plasmids (∼61 and 67 kbp) were not recovered in the resulting assembly. These plasmids were visible in the Eckhardt-type gel (protocol I, see above; Fig. S1A). One of these two plasmids was also missing when the best 2 Gbp of reads (>6 kbp read length) were used for assembly with Flye. This is not unexpected when using super-high-depth read sets with Filtlong, since the resulting subsets are enriched in longer reads and depleted in shorter ones, leading to the loss of some smaller plasmids during assembly due to insufficient coverage or representation. Therefore, the “--meta” option was additionally applied, which enabled recovery of both small circular plasmids. Interestingly, in three of the four Flye assemblies, an ∼80 kbp linear contig was recovered. The only exception was the Flye assembly based on the best 1 Gbp of reads without the --meta flag, from which this linear replicon was absent. The other long-read assemblers we tested (Miniasm/Minipolish and Canu) also failed to recover this replicon, except for Raven when using the best 2 Gbp of reads as input, although the ∼80 kbp contig was assigned as circular in this assembly. On the other hand, assemblies generated with software that used both long and short reads (Hybracter, Unicycler and Spades) as an input did include this ∼80 kbp contig. Assembly with Hybracter and Unicycler included a circularization step, but as for the Flye assemblies, an ∼80 kbp contig was assigned as linear. Therefore, the authenticity and topology of this replicon in rho-13.3 was further assessed.

**Table 2.** General features of the genome sequences obtained in this study.

|  | Strains |  |  |  |  |  |
| --- | --- | --- | --- | --- | --- | --- |
|  | *Allorhizobium*<br>sp. Av2 | *Agrobacterium*<br>*rosae* rho-7.1 | *Agrobacterium*<br>sp. rho-8.1 | *Agrobacterium*<br>*rosae* rho-11.1 | *Agrobacterium*<br>sp. rho-13.3 | *Agrobacterium*<br>*rosae* rho-14.1 |
| Replicons (N) | 6 | 7 | 9 | 7 | 7 | 10 |
| Size (Mbp) | 6.34 | 5.65 | 5.55 | 6.03 | 5.76 | 5.99 |
| GC content (%) | 57.61 | 56.55 | 56.68 | 56.64 | 56.17 | 56.57 |
| Genes<sup>a</sup> (total; N) | 5,812 | 5,402 | 5,316 | 5,756 | 5,461 | 5,729 |
| CDSs<sup>a</sup> (with protein; N) | 5,630 | 5,237 | 5,123 | 5,543 | 5,307 | 5,537 |
| 5S-23S-16S rRNA operons (N) | 4 | 4 | 4 | 4 | 4 | 4 |
| Mean depth of genome<br>coverage (x)<sup>b</sup> | 353.09 | 487.12 | 602.07 | 379.48 | 1811.44 | 399.33 |
| Accession number |  |  |  |  |  |  |

<sup>a</sup>Numbers based on PGAP annotation.
<sup>b</sup>Mean depth of genome coverage based on both Nanopore and Illumina data.

First, a rho-13.3 genome assembly using only short reads was performed with Spades (with option “--isolate”) and assembly graphs were examined. Interestingly, the ∼80- kbp contig corresponding to the putative linear replicon was connected at both ends to the same end of another 1,096-bp contig. This two-contig structure is a signature of linear plasmids carrying terminal inverted repeats (TIRs) [18]. In other words, the assembly graph suggested that the shorter 1,096-bp contig represents a TIR at both ends of the ∼80-kbp contig. Consistent with this interpretation, the depth of coverage for this short contig (as indicated by Spades), was double that of the longer contig. This putative linear replicon was then manually assembled and the long Nanopore reads (>30 kbp) were mapped to the final assembly. The read alignment examination clearly showed that numerous long reads soft-clipped over the edges of the plasmid sequence. Further examination showed that these reads continue past the end into the other strand, evidently indicating that the plasmid ends have a hairpin structure with covalently closed ends. This allowed us to identify the point where the loop occurs and the definite end of the linear plasmid. Similarly, the hairpin ends of the linear chromid, which is also carried by rho-13.3, were identified and the replicon sequence was manually finished.

To identify if similar plasmids occur in other bacterial strains with publicly available genomes, blastp searches against the NCBI non-redundant protein sequence (nr) database were performed. Not surprisingly, nonpathogenic *Agrobacterium* strains sequenced in our previous study (rho-7.1, rho-8.1, rho-11.1 and rho-14.1) [21], which were also isolated from aerial crown gall tumors on rhododendron from the same locality in Germany as strain rho- 13.3, were the best hits. Surprisingly, the next-best hit was *Allorhizobium* sp. Av2, which was sequenced in a separate study from our group [6], but was isolated from grapevine crown gall originating from Croatia. As only draft genomes were available for these additional strains, we performed ONT sequencing to generate their complete genome sequences. For strains Av2, rho-7.1, rho-8.1, rho-11.1, and rho-14.1, whole-genome ONT sequencing generated 1.2-2.4 Gbp of data per bacterial strain, enabling a high depth of genome coverage (>200×; Table S1). To improve the read length N50 for genome assembly, reads less than 6 kbp in length and 10% of the worst reads (low accuracy) were discarded, resulting in read length N50 values >15 kbp and depths of genome coverage >120× (Table S1). As is typical for *Agrobacterium* spp., genome assemblies of strains rho-7.1, rho-8.1, rho- 11.1, and rho-14.1 included a large circular chromosome, a linear chromid, and multiple plasmids, with the total number of replicons ranging from 6 to 10 (Table 2). On the other hand, in addition to the circular chromosome, *Allorhizobium* sp. Av2 carried a circular chromid and multiple plasmids. Among the plasmids, the *Agrobacterium* strains carried an opine-catabolic (OC) plasmid characterized before [21], while the strain Av2 carried a tumor-inducing (Ti) plasmid required for pathogenicity. Following analyses similar to those reported above for rho-13.3, the resulting genome assemblies confirmed the presence of a linear plasmid in all the analyzed strains.

Interestingly, *Agrobacterium rosae* strain rho-6.1, whose complete genome sequence was reported before by our group and which originated from the same locality in Germany as the other “rho” strains sequenced in the current study, appeared not to carry a linear plasmid. This strain was previously sequenced using PacBio and Illumina approaches [21]. It was assembled using a long-read first approach using the “RS_HGAP_Assembly.3” protocol included in SMRT Portal version 2.3.0, followed by error-correction with Illumina reads. In the current study, we additionally inspected an Illumina-only Spades assembly and assembly graphs, which was also consistent with the linear plasmid being absent from this strain. Moreover, we designed primers targeting a putatively conserved gene of the linear plasmids revealed in this study, and performed PCR analysis. However, specific amplification was not observed, confirming that the strain rho-6.1 does not carry a linear plasmid.

### Experimental validation of the presence of linear plasmids in the family *Rhizobiaceae*

In order to validate the genome assemblies, primarily the authenticity of the linear plasmid, Eckhardt-type gel electrophoresis of complete genomic DNA was conducted using two protocols. Protocol I was tailored for separation of circular plasmids and involves a longer electrophoresis step (4 V cm^-1^ for 20 h) [4]. It allowed clear separation of the circular plasmids (61, 67, 155 and 438 kbp) of strain rho-13.3 (Fig. S1A). However, the replicon corresponding to the linear plasmid was not visible in this Eckhardt-type gel (Fig. S1A). Therefore, another slightly different Eckhardt-type protocol (protocol II) was applied, which involves a shorter electrophoretic step (3.2 V/cm for 6-10 h). This time, two additional bands were resolved for strain rho-13.3, showing faster migration speed compared to the circular plasmids (Fig. S1B). These bands may have migrated out of the end of gel when protocol I was used. In any case, we assumed that these two bands might correspond to the linear replicons carried by this strain, the larger one being the linear chromid, and the smaller one being the linear plasmid (Fig. S1B). Likewise, a band most likely corresponding to the linear plasmid was also resolved for *Allorhizobium* sp. Av2 (Fig. S1B). *Allorhizobium* sp. Av2 does not carry a linear chromid, unlike Agrobacterium spp. (i.e., rho-13.3), and thus as expected, only one band migrating faster than the circular replicons was visualized in the gel for the strain Av2 (Fig. S1B).

To estimate the size of the bands assumed to correspond to linear plasmids of strains rho-13.3 and Av2, the bands were compared to a Quick-Load 1 kb Extend DNA Ladder (New England BioLabs), which includes digested, linear DNA fragments ranging from 0.5 to 48.5 kbp. The linear plasmid-bands migrated slightly slower than the largest 48.5-kbp band of the DNA ladder (Fig. S1B). Interestingly, the linear plasmids of strains rho-13.3 and Av2, with sizes of 81 and 79 kbp, respectively, migrated faster than the circular 79-kbp plasmid of the strain All. ampelinum S4 (Fig. S1B). The linear 79-kbp plasmid even migrated faster than the circular 44-kbp plasmid of the reference strain R. rhizogenes K84 (Fig. S1C). In other words, putative linear plasmids revealed by sequencing migrated faster than expected on Eckhardt-type gels. For instance, when comparing different topological forms of the same plasmid (e.g., pUC19 of size 2.7 kbp), a supercoiled circular form migrated the fastest, followed by a linear and open circular (oc; relaxed structure) form [59], However, the migration speed of different plasmid forms can change with the electrophoresis conditions, such as agarose concentration, electric-field strength and ionic conditions [60,61]. Nevertheless, the standard agarose gel electrophoresis of purified DNA was used in these examples from the literature, which most likely can affect supercoiling of replicons and their mobility in the gel, compared to Eckhardt-type gel electrophoresis relying on electrophoresis of bacterial cells that are lysed directly in the gel.

Taken together, Eckhardt-type gel electrophoresis suggested that the fast-migrating bands of strains rho-13.3 and Av2 might indeed correspond to the linear plasmids revealed by whole-genome sequencing. However, these data alone could not rule out that the fast-migrating bands were not either fragments of sheared DNA or circular plasmids in a compact supercoiled form. Therefore, to further verify the assignments of the fast-migrating bands as linear plasmids, they were cut from the gel and DNA was purified. Bands thought to correspond to the linear plasmids of strains Agrobacterium sp. rho-13.3 and Allorhizobium sp. Av2 were cut from two independent Eckhardt-type gel and subjected to Illumina high-throughput sequencing. The vast majority of the reads mapped back to the linear plasmid, while a small minority of reads mapped to other replicons (Fig. 1; Table S2). For example, for the band thought to correspond to the linear plasmid (pLin13.3) of strain rho-13.3, when the sequencing reads were mapped to the rho-13.3 genome, the mean depth of coverage exceeded 25,000× for pLin13.3, while it ranged from ∼10-110× for the other replicons (Fig. 1; Table S2). The detection of a minority of DNA from other replicons is not unexpected as sheared DNA of other replicons may comigrate with the DNA of an individual replicon [62,63]. Taken together, the sequencing results strongly indicate that the fast-migrating bands indeed represent linear plasmids.

**Figure 1.**
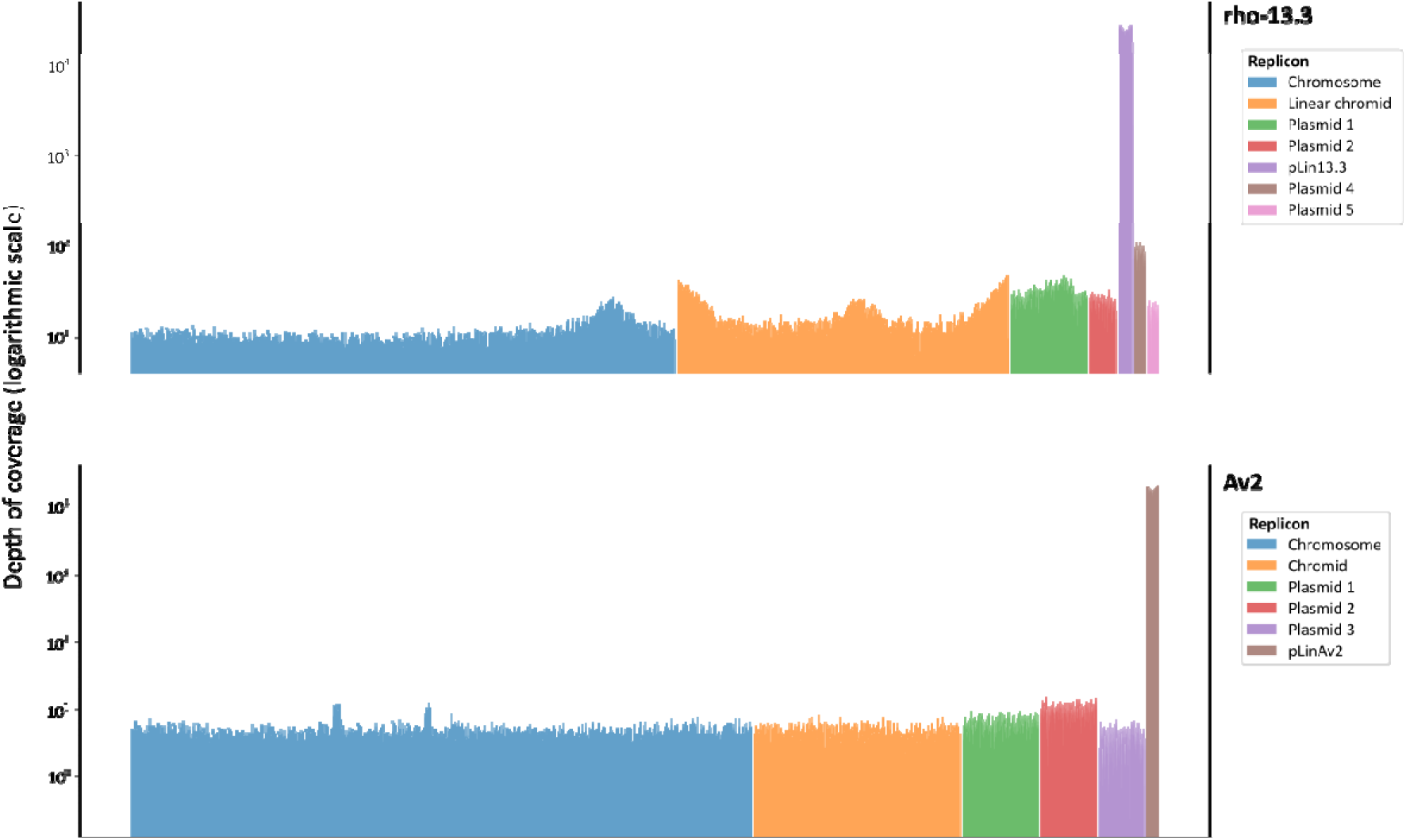
Depth of coverage along replicons. Bands corresponding to linear plasmids of strains Agrobacterium sp. rho-13.3 and Allorhizobium sp. Av2 were cut from the Eckhardt- type gel, DNA was purified and sequenced using Illumina platform (see Materials and Methods for more detail). Plots show results for one representative DNA sample per strain (Table S2). Bars correspond to the mean coverage of 5,000 nucleotide windows for the chromosomes, 2,000 nucleotide windows for chromids, 500 nucleotide windows for plasmid 1 of both strains, 200 nucleotide windows for plasmid 2 (rho-13.3) and plasmids 2 and 3 (Av2), and 100 nucleotide windows for pLin13.3, pLinAv2, and plasmids 4 and 5 (rho-13.3).

Moreover, the band presumably corresponding to the linear chromid of the strain rho-13.3 was also purified from the gel and sequenced. For this DNA sample, the mean depth of coverage reached 801× for the linear chromid, while it was <50× for other replicons, except for pLin13.3 for which it reached 233× (Fig. S2; Table S2). This band indeed seems to primarily correspond to the linear chromid DNA (most likely sheared), although it contained relatively high amount of DNA corresponding to the linear plasmid (pLin13.3), which was resolved in the close proximity in the Eckhardt-type gel (Fig. S1B).

PFGE was used to further validate the presence of linear plasmids in the six sequenced strains. For this purpose, different pulse times were used, enabling separation of either large or small replicons. By using pulse time of 10-40 s, it was possible to separate all plasmids of strains rho-13.3 and Av2, including the linear plasmid (Fig. S3A). Unlike Eckhardt-type gel electrophoresis, in PFGE, topology (linear versus circular) tends not to influence the mobility of a replicon (Fig. S3). For instance, a band presumed to correspond to the pLin13.3 migrated slower than a band corresponding to the small circular plasmids (61 and 67 kbp) of strain rho-13.3 (Fig. S3A). As these two small circular plasmids could not be resolved by using pulse time of 10-40 s, we confirmed the presence of these two plasmids by using pulse time of 1-25 s, which is more suitable for separation of smaller replicons (Fig. S3B).

Moreover, PFGE was also performed with plugs treated with S1 nuclease. S1 nuclease targets single-stranded DNA that can occur in negatively supercoiled plasmids, and converts them into the linear form [64,65]. In this respect, it has been reported that the circular and linear forms of plasmids can be distinguished because the supercoiled circular forms migrate more slowly than their corresponding linear forms [65,66]. Surprisingly, under the PFGE conditions used in this study, treated replicons migrated slightly slower than those treated with S1 nuclease (Fig. S3C). In the case of circular plasmids, S1 nuclease treatment might result in less compact, slower-migrating forms. However, it was interesting that a similar delay in mobility was also evident for the linear plasmid of the strain rho-13.3 (Fig. S3C). Linear plasmids identified in this study may be similar to the linear plasmids of Borrelia burgdorferi, which have covalently closed ends and contain a single-stranded loop at each end [67]. This plasmid form can be structurally more stable and compact, migrating more efficiently in the gel. An S1 nuclease might cleave the hairpin ends, and likely introduce additional nicks, which could cause the linear plasmid to become less compact and to have slower migration in the PFGE gel.

### Genomic comparison of the linear plasmids

The GC content of the linear plasmids sequenced in this study ranged from 53.73 to 53.93%, which is lower than the GC content of the chromosomes in these same strains, which range from 56.43 to 57.02% for the five Agrobacterium strains and 57.89% for *Allorhizobium* sp. Av2 (Tables 2 and 3). The substantial difference in GC content between the linear plasmids and the chromosomes suggest that the linear plasmids were likely acquired more recently through horizontal gene transfer.

The size of the linear plasmids ranged from 76,871 (pLin8.1) to 83,153 bp (pLin11.1) (Table 3). The differences in the size were primarily due to insertion sequence (IS) elements present in some plasmids (Fig. 2). Plasmids pLin7.1 and pLin14.1 were identical, and they differed from pLin13.3 in only one single nucleotide polymorphism (SNP). Compared to these three plasmids, plasmid pLin11.1 possessed a two additional IS elements (Fig. 2). On the other hand, plasmid pLin8.1 had one fewer IS element compared to pLin7.1, pLin13.3, and pLin14.1, but contained a separate additional 2571-bp fragment (Fig. 2). Compared to these five plasmids identified in the Agrobacterium strains, pLinAv2 carried by *Allorhizobium* sp. Av2 has several variations. In particular, a region including a parB gene, an IS3 element, a toxin-antitoxin (TA) system, and several genes encoding hypothetical proteins (HPs) were absent in this plasmid, but unlike the other plasmids, it carried different region comprising a distinct TA system as well as several genes coding for HPs (Fig. 2). Moreover, the right end of plasmid pLinAv2 showed a slightly lower degree of homology to the plasmids carried by the *Agrobacterium* strains (Fig. 2). Overall, despite the differences described above, all six linear plasmids analyzed showed a high degree of homology, clearly indicating their common ancestry.

**Fig. 2.**
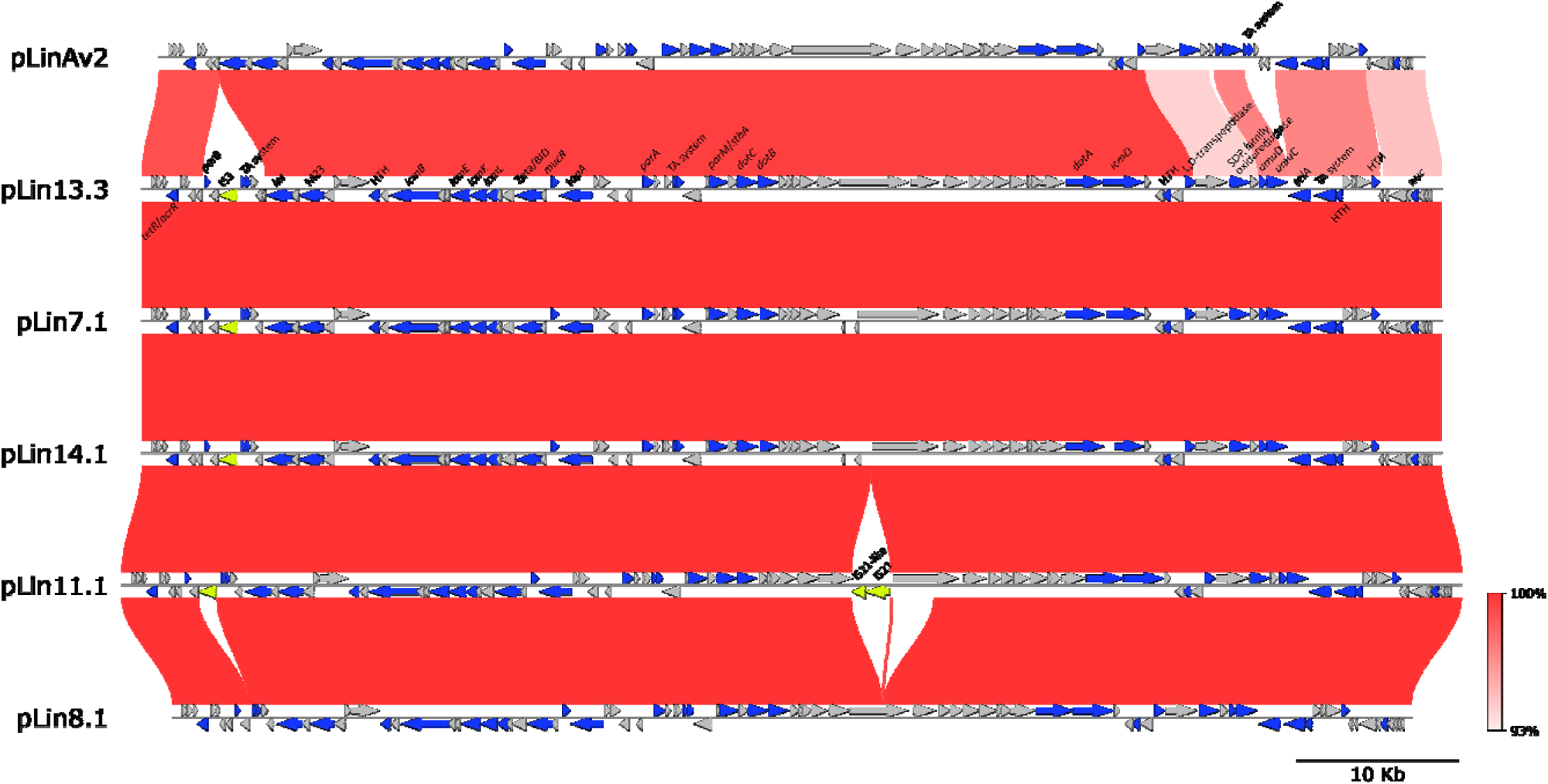
Synteny and comparative analysis of linear plasmids sequenced in this study. MUMmer alignment and visualization were performed with ‘pgv- mummer’ workflow, which is part of the python genome visualization package ‘pyGenomeViz’ version 1.5.0 (https://github.com/moshi4/pyGenomeViz). The colored arrows represent coding sequences (CDSs): annotated genes (blue arrows), insertion sequence (IS) elements (yellow arrows) and genes encoding hypothetical proteins (grey arrows). Gene names are indicated above the arrows only for the reference plasmid pLin13.3 (for more information see Table S3) and for other plasmids only for the genes that are absent from pLin13.3. The red blocks connecting different gene regions of two plasmids indicate the identity between them. The darker color indicates a higher percentage of identity.

**Table 3.**
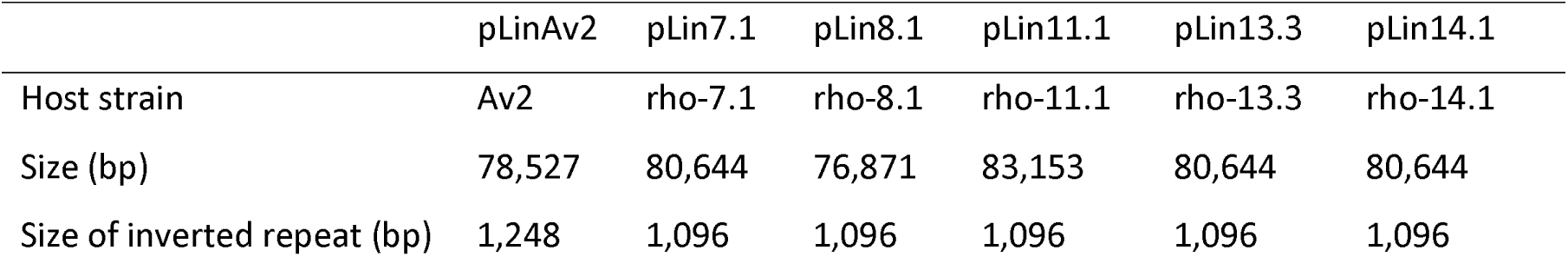
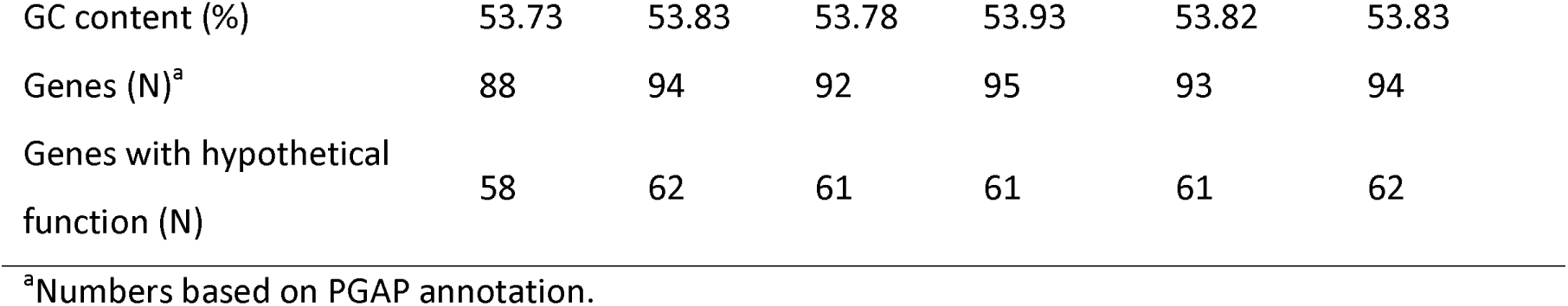
General features of linear plasmids investigated in this study.

### Gene content and functional annotation of linear plasmids

The putative functions of the majority of genes carried by the linear plasmids remain unknown, and they were annotated as encoding hypothetical proteins (Table 3). In addition to the hypothetical genes, all the linear plasmids (Table 3) carried a gene putatively encoding a protelomerase (Table S3; Fig. 2), further supporting that they have a linear topology with closed hairpin ends. Unlike *Allorhizobium* sp. Av2, which does not carry additional linear replicons, the *Agrobacterium* spp. strains (Table 2) carried another gene putatively encoding a protelomerase on the linear chromids. A gene encoding a protelomerase was located on the linear chromids for some other members of the Agrobacterium sub-clade “rubi” for which complete genome assemblies were available, i.e. Agrobacterium vaccinii B7.6^T^ (CP054151). This is also true for Agrobacterium larrymoorei CFBP 5477 (CP124734). Interestingly, in members of the Agrobacterium “biovar 1” sub- clade, such as the strains C58^T^ and H13-3, it was reported that a protelomerase gene responsible for chromid linearity is encoded on their circular chromosome rather than on the chromid itself [13,14,68]. Protelomerases of the linear plasmids were distinct from those associated with the linear chromids of Agrobacterium spp. For instance, protelomerase of pLin13.3 shared 67.3% amino acid identity with protelomerases associated with linear chromids of the same strain rho-13.3.

The linear plasmids harbored genes putatively involved in plasmid partitioning (parA, parM/stbA) and multiple TA systems putatively associated with plasmid maintenance, although they may also play roles in stress responses or phage resistance (Table S3; Fig. 2). Other genes potentially associated with stress tolerance or DNA damage include genes putatively encoding some components of DNA polymerase V (subunits C and D). Interestingly, DNA polymerase V is an error-prone polymerase [69], and some rhizobia encode error-prone DNA polymerases that appear to result in elevated mutation rates during legume symbiosis [70]. It is therefore tempting to speculate that the DNA polymerase V encoded by the linear plasmids may similarly result in elevated mutation rates during tumorigenesis. In addition, all six linear plasmids carried genes putatively encoding an apolipoprotein N-acyltransferase (lnt), a M23 family metallopeptidase (M23), and a L,D-transpeptidase (Table S3; Fig. 2) involved in lipoprotein production, lysis of bacterial cell wall peptidoglycans, and in peptidoglycan cross-linking, respectively. These functions might have a role in cell wall remodeling for persistence or survival under stress. A gene putatively coding for a type I DNA topoisomerase carried by these plasmids is most likely involved in relieving supercoiling of linear DNA during DNA replication and transcription (Table S3; Fig. 2).

Moreover, dot and icm genes putatively associated with the type IV secretion system (T4SS) were conserved in all six linear plasmids (Table S3; Fig. 2). In general, dot/icm genes encode a T4SS classified as type IVB (T4SS^Dot/Icm^) which is evolutionarily most closely related to conjugation systems of IncI plasmids, while a distinct type IVA resembles the VirB/VirD4 T4SS of *Agrobacterium tumefaciens* [71,72]. The T4SS^Dot/Icm^ is present in intracellular bacterial pathogens such as in *Legionella pneumophila* and related bacteria, where it is crucial for virulence by translocating a large number of effectors into the eukaryotic host cell [73]. However, dot/icm gene clusters were also identified in diverse bacteria, including the genera *Burkholderia, Methylorubrum,* and *Xanthomonas*, where they were hypothesized to play a role in conjugation [74]. Indeed, it was demonstrated that the T4SS^Dot/Icm^ of L. pneumophila has the capacity to transfer plasmid DNA from one cell to another [75]. Additional BLAST searches also revealed some homology (<53% of identity for ∼50% query cover) between a gene located downstream the *icmL* gene (Zeta/BID; Fig. 2) and proteins annotated as “zeta toxin family proteins” as well as “BID domain-containing T4SS effectors” (Table S3). Interestingly, the BID domain is present in alphaproteobacterial toxins, relaxases, and *Bartonella* effector proteins [76]. Taken together, it is tempting to speculate that the T4SS^Dot/Icm^ identified in the linear plasmids are involved in their conjugation, although we could not identify a corresponding relaxase gene or an oriT. Alternatively, considering the other described functions of T4SSs [77], it might play a role in bacterial competition by delivering toxins to competing bacteria, or by injecting effector proteins to plant cells. Nevertheless, apart from the Zeta/BID gene whose function is highly speculative, we could not identify type IV effector proteins in the linear plasmids.

### The distribution and evolution of linear plasmids

To examine whether additional organisms carried linear plasmids related to the six linear plasmids identified as part of this study, we searched GenBank using the protein sequences of the telomere resolvase and DNA topoisomerase I as queries against the nr protein database. This led to the identification of an additional six replicons/contigs that carried both a telomere resolvase and a DNA topoisomerase I, and for which raw sequencing data were available, allowing further examination (Table S4). The corresponding genes of these six newly identified strains shared low sequence identity with the telomere resolvase (∼28- 30%) and DNA topoisomerase I (∼42-44%) proteins of the six linear plasmids previously identified in this study. All six strains belonged to the family *Rhizobiaceae*, including one *Allorhizobium* strain, three *Rhizobium* strains, and two *Agrobacterium* strains (Table S4). For the four strains for which short-read data were available (Table S4), assembly using Spades and/or Shovil was performed, and assembly graphs were examined. In all cases, the replicons carrying a telomere resolvase and a DNA topoisomerase I were connected at both ends to the same end of another short contig, a signature of plasmids with a linear topology. For the other two strains, for which only long-read data was available, we inspected the corresponding contigs and identified hairpin ends (Table S4). Overall, these results are consistent with these six new *Rhizobiaceae* replicons also representing linear plasmids.

Whole-plasmid AAI comparisons showed that the six putatively linear plasmids discovered by GenBank mining are diverse (Table S5, Fig. S4). Nevertheless, putative linear plasmids of strains AB3 and AC44/96 were relatively closely related (AAI 91.6%), as were the putatively linear plasmids of strains NCPPB 2655, CECT 9324, and T5/73 (AAI ≥82.7%). The six newly identified plasmids also carried *dot*/*icm* genes encoding a T4SS. Interestingly, the putative linear plasmid of strain AB3 carried a gene annotated as a relaxase/mobilization nuclease domain-containing protein (GenBank Acc. No. NZ_MAVS02000024.1, locus_tag BBI12_024275). In the future, it would be interesting to obtain finished sequences of these six new putatively linear plasmids.

As suggested by the single gene comparisons mentioned above, whole-plasmid AAI analyses confirmed that the six new, putatively linear plasmids were distantly related to the linear plasmids identified earlier in our study (Table S5, Fig. S4). This indicates that these linear plasmids have independent evolutionary histories. Similarly, these two groups of linear plasmids formed well-separated clusters in a phylogenetic tree based on protelomerase protein sequences (Fig. S5). Interestingly, protelomerases associated with the linear plasmids sequenced in this study formed a monophyletic group intertwined within the protelomerases of the linear chromids (Fig. S5), potentially suggesting that they evolved from a protelomerase acquired from the agrobacterial linear chromids.

## 8. Conclusion

Overall, our results are consistent with linear plasmids being acquired multiple independent times by diverse members of the family *Rhizobiaceae*. To the best of our knowledge, linear plasmids have not previously been identified in the family *Rhizobiaceae* or the class *Alphaproteobacteria* to which the family *Rhizobiaceae* belongs. The identification of linear plasmids in members of the family *Rhizobiaceae* provides further evidence of their extraordinary genome plasticity, and expands the taxonomic range in which linear plasmids have been identified. Linear plasmids were identified in three *Rhizobiaceae* genera, and they were found even in published genome assemblies that were not previously recognized to have linear plasmids. These results suggest that linear plasmids may be even more widespread in the family *Rhizobiaceae* than what was detected in this study, and that they may go undetected in genome sequencing studies if the assemblies are not specifically examined for linear plasmid. The linear plasmids identified in this study are carried by both nonpathogenic and tumorigenic organisms, although their biological functions remain unknown. In the future, it will be interesting to further explore the roles of these plasmids in the biology of these organisms.

## 9 Author statements

### 9.1 Author contributions

Conceptualization: N.K. Data Curation: N.K., R.R.W., M.K., G.C.D. and T.T.. Formal Analysis: N.K., R.R.W., M.K., G.C.D., J.P., M.F.H.. Funding Acquisition: N.K., K.S., J.P. and D.B.

Investigation: N.K. and M.K. Methodology: N.K., M.K., J.P. and M.F.H. Resources: N.K., K.S., T.T. and D.B. Supervision: N.K. Validation: N.K. and R.R.W. Visualization: N.K. and M.K. Writing – Original Draft: N.K. Writing – Review & Editing: all authors.

### 9.4 Ethical approval

Not applicable.

### 9.4 Consent for publication Not applicable

## Acknowledgments

This research was enabled, in part, through computational resources provided by BMBF-funded de.NBI Cloud within the German Network for Bioinformatics Infrastructure (de.NBI) (031A532B, 031A533A, 031A533B, 031A534A, 031A535A, 031A537A, 031A537B, 031A537C, 031A537D, 031A538A). We thank Ute Zimmerling (JKI-EP) for excellent technical assistance.

## References

[1] diCenzo GC, Yang Y, Young JPW, Kuzmanović N. Refining the taxonomy of the order *Hyphomicrobiales* (*Rhizobiales*) based on whole genome comparisons of over 130 type strains, Int J Syst Evol Microbiol 2024;74.

[2] Carareto Alves LM, Souza JAM de, Varani AdM, Lemos, Eliana Gertrudes de Macedo. The family *Rhizobiaceae*. In: Rosenberg E, DeLong EF, Lory S, Stackebrandt E, Thompson F (editors). The Prokaryotes: Alphaproteobacteria and Betaproteobacteria. Berlin, Heidelberg: Springer Berlin Heidelberg; 2014. pp. 419–437.

[3] de Lajudie PM, Andrews M, Ardley J, Eardly B, Jumas-Bilak E et al. Minimal standards for the description of new genera and species of rhizobia and agrobacteria, Int J Syst Evol Microbiol 2019;69:1852–1863.

[4] Kuzmanović N, diCenzo GC, Bunk B, Spröer C, Frühling A et al. Genomics of the “tumorigenes” clade of the family *Rhizobiaceae* and description of *Rhizobium rhododendri* sp. nov, MicrobiologyOpen 2023;12:e1352.

[5] Escobar MA, Dandekar AM. *Agrobacterium tumefaciens* as an agent of disease, Trends in plant science 2003;8:380–386.

[6] Kuzmanović N, Biondi E, Overmann J, Puławska J, Verbarg S et al. Genomic analysis provides novel insights into diversification and taxonomy of *Allorhizobium vitis* (i.e. *Agrobacterium vitis*), BMC genomics 2022;23:462.

[7] Geddes BA, Kearsley J, Morton R, diCenzo GC, Finan TM. Chapter Eight - The genomes of rhizobia. In: Frendo P, Frugier F, Masson-Boivin C (editors). Advances in Botanical Research. Academic Press; 2020. pp. 213–249.

[8] diCenzo GC, Finan TM. The divided bacterial genome: structure, function, and evolution, Microbiol Mol Biol Rev 2017;81.

[9] Harrison PW, Lower RP, Kim NK, Young JP. Introducing the bacterial ’chromid’: not a chromosome, not a plasmid, Trends Microbiol 2010;18:141–148.

[10] Hall JPJ, Botelho J, Cazares A, Baltrus DA. What makes a megaplasmid?, Philosophical Transactions of the Royal Society B: Biological Sciences 2022;377:20200472.

[11] Allardet-Servent A, Michaux-Charachon S, Jumas-Bilak E, Karayan L, Ramuz M. Presence of one linear and one circular chromosome in the *Agrobacterium tumefaciens* C58 genome, J Bacteriol 1993;175:7869–7874.

[12] Ramírez-Bahena MH, Vial L, Lassalle F, Diel B, Chapulliot D et al. Single acquisition of protelomerase gave rise to speciation of a large and diverse clade within the *Agrobacterium*/*Rhizobium* supercluster characterized by the presence of a linear chromid, Mol Phylogenet Evol 2014;73:202–207.

[13] Slater S, Setubal JC, Goodner B, Houmiel K, Sun J et al. Reconciliation of sequence data and updated annotation of the genome of *Agrobacterium tumefaciens* C58, and distribution of a linear chromosome in the genus *Agrobacterium*, Appl Environ Microbiol 2013;79:1414–1417.

[14] Huang WM, DaGloria J, Fox H, Ruan Q, Tillou J et al. Linear chromosome-generating system of Agrobacterium tumefaciens C58: protelomerase generates and protects hairpin ends, J Biol Chem 2012;287:25551–25563.

[15] Naranjo HD, Lebbe L, Cnockaert M, Lassalle F, Too CC et al. Phylogenomics reveals insights into the functional evolution of the genus *Agrobacterium* and enables the description of *Agrobacterium divergens* sp. nov, Syst Appl Microbiol 2023;46:126420.

[16] Kinashi H. Linear Plasmids from Actinomycetes, Nippon Hosenkin Gakkaishi 1994;8:87– 96.

[17] Chaconas G, Kobryn K. Structure, function, and evolution of linear replicons in Borrelia, Annu Rev Microbiol 2010;64:185–202.

[18] Hawkey J, Cottingham H, Tokolyi A, Wick RR, Judd LM et al. Linear plasmids in Klebsiella and other Enterobacteriaceae, Microb Genom 2022;8.

[19] Baker S, Hardy J, Sanderson KE, Quail M, Goodhead I et al. A novel linear plasmid mediates flagellar variation in Salmonella Typhi, PLoS Pathog 2007;3:e59.

[20] Kuzmanović N, Biondi E, Bertaccini A, Obradović A. Genetic relatedness and recombination analysis of *Allorhizobium vitis* strains associated with grapevine crown gall outbreaks in Europe, Journal of applied microbiology 2015;119:786–796.

[21] Kuzmanović N, Nesme J, Wolf J, Neumann-Schaal M, Petersen J et al. Deciphering the key players within the bacterial microbiota associated with aerial crown gall tumors on rhododendron: Insights into the gallobiome, Phytobiomes J 2024;8:401–415.

[22] Danecek P, Bonfield JK, Liddle J, Marshall J, Ohan V et al. Twelve years of SAMtools and BCFtools, GigaScience 2021;10.

[23] Wick RR, Judd LM, Holt KE. Assembling the perfect bacterial genome using Oxford Nanopore and Illumina sequencing, PLoS Comput Biol 2023;19:e1010905.

[24] Kolmogorov M, Yuan J, Lin Y, Pevzner PA. Assembly of long, error-prone reads using repeat graphs, Nature Biotechnology 2019;37:540–546.

[25] Kosakovsky Pond SL, Posada D, Gravenor MB, Woelk CH, Frost SD. GARD: a genetic algorithm for recombination detection, *Bioinformatics (Oxford*, England) 2006;22:3096– 3098.

[26] Wick RR, Holt KE. Benchmarking of long-read assemblers for prokaryote whole genome sequencing, F1000Res 2019;8:2138.

[27] Li H. Minimap and miniasm: fast mapping and de novo assembly for noisy long sequences, Bioinformatics 2016;32:2103–2110.

[28] Vaser R, Sović I, Nagarajan N, Šikić M. Fast and accurate de novo genome assembly from long uncorrected reads, Genome Res 2017;27:737–746.

[29] Koren S, Walenz BP, Berlin K, Miller JR, Bergman NH et al. Canu: scalable and accurate long-read assembly via adaptive k-mer weighting and repeat separation, Genome Research 2017;27:722–736.

[30] Wick RR, Judd LM, Cerdeira LT, Hawkey J, Méric G et al. Trycycler: consensus long-read assemblies for bacterial genomes, Genome Biology 2021;22:266.

[31] Wick RR, Howden BP, Stinear TP. Autocycler: long-read consensus assembly for bacterial genomes, bioRxiv 2025:2025.05.12.653612.

[32] Wick RR, Holt KE. Polypolish: Short-read polishing of long-read bacterial genome assemblies, PLoS Comput Biol 2022;18:e1009802.

[33] Bouras G, Houtak G, Wick RR, Mallawaarachchi V, Roach MJ et al. Hybracter: enabling scalable, automated, complete and accurate bacterial genome assemblies, Microb Genom 2024;10.

[34] Prjibelski A, Antipov D, Meleshko D, Lapidus A, Korobeynikov A. Using SPAdes de novo assembler, Current protocols in bioinformatics 2020;70:e102.

[35] Li H. Minimap2: pairwise alignment for nucleotide sequences, Bioinformatics (Oxford, England) 2018;34:3094–3100.

[36] Li H, Durbin R. Fast and accurate short read alignment with Burrows-Wheeler transform, *Bioinformatics (Oxford*, England) 2009;25:1754–1760.

[37] Wick RR, Schultz MB, Zobel J, Holt KE. Bandage: interactive visualization of de novo genome assemblies, Bioinformatics 2015;31:3350–3352.

[38] Tatusova T, DiCuccio M, Badretdin A, Chetvernin V, Nawrocki EP et al. NCBI prokaryotic genome annotation pipeline, Nucleic Acids Res 2016;44:6614–6624.

[39] Cantalapiedra CP, Hernández-Plaza A, Letunic I, Bork P, Huerta-Cepas J. eggNOG- mapper v2: functional annotation, orthology assignments, and domain prediction at the metagenomic scale, Mol Biol Evol 2021;38:5825–5829.

[40] Huerta-Cepas J, Szklarczyk D, Heller D, Hernández-Plaza A, Forslund SK et al. eggNOG 5.0: a hierarchical, functionally and phylogenetically annotated orthology resource based on 5090 organisms and 2502 viruses, Nucleic Acids Res 2018;47:D309–D314.

[41] Buchfink B, Reuter K, Drost H-G. Sensitive protein alignments at tree-of-life scale using DIAMOND, Nat Methods 2021;18:366–368.

[42] Johnson M, Zaretskaya I, Raytselis Y, Merezhuk Y, McGinnis S et al. NCBI BLAST: a better web interface, Nucleic Acids Res 2008;36:W5–W9.

[43] Kurtz S, Phillippy A, Delcher AL, Smoot M, Shumway M et al. Versatile and open software for comparing large genomes, Genome biology 2004;5.

[44] Kim D, Park S, Chun J. Introducing EzAAI: a pipeline for high throughput calculations of prokaryotic average amino acid identity, J Microbiol 2021;59:476–480.

[45] Hyatt D, Chen G-L, Locascio PF, Land ML, Larimer FW et al. Prodigal: prokaryotic gene recognition and translation initiation site identification, BMC Bioinformatics 2010;11:119.

[46] Steinegger M, Söding J. MMseqs2 enables sensitive protein sequence searching for the analysis of massive data sets, Nature Biotechnology 2017;35:1026–1028.

[47] Katoh K, Rozewicki J, Yamada KD. MAFFT online service: multiple sequence alignment, interactive sequence choice and visualization, Briefings Bioinf 2019;20:1160–1166.

[48] Nguyen L-T, Schmidt HA, Haeseler A von, Minh BQ. IQ-TREE: a fast and effective stochastic algorithm for estimating maximum-likelihood phylogenies, Mol Biol Evol 2015;32:268–274.

[49] Trifinopoulos J, Nguyen L-T, von Haeseler A, Minh BQ. W-IQ-TREE: a fast online phylogenetic tool for maximum likelihood analysis, Nucleic Acids Res 2016;44:W232–W235.

[50] Kalyaanamoorthy S, Minh BQ, Wong TKF, Haeseler A von, Jermiin LS. ModelFinder: fast model selection for accurate phylogenetic estimates, Nat Methods 2017;14:587– 589.

[51] Schwarz G. Estimating the dimension of a model, The Annals of Statistics 1978;6:461–464, 4.

[52] Hoang DT, Chernomor O, Haeseler A von, Minh BQ, Le Vinh S. UFBoot2: improving the ultrafast bootstrap approximation, Mol Biol Evol 2018;35:518–522.

[53] Guindon S, Dufayard JF, Lefort V, Anisimova M, Hordijk W et al. New algorithms and methods to estimate maximum-likelihood phylogenies: assessing the performance of PhyML 3.0, Systematic biology 2010;59:307–321.

[54] Untergasser A, Cutcutache I, Koressaar T, Ye J, Faircloth BC et al. Primer3--new capabilities and interfaces, Nucleic acids Res 2012;40:e115.

[55] Kuzmanović N, Prokić A, Ivanović M, Zlatković N, Gašić K et al. Genetic diversity of tumorigenic bacteria associated with crown gall disease of raspberry in Serbia, Eur J Plant Pathol 2015;142:701–713.

[56] Eckhardt T. A rapid method for the identification of plasmid desoxyribonucleic acid in bacteria, Plasmid 1978;1:584–588.

[57] Hynes MF, Simon R, Pühler A. The development of plasmid-free strains of Agrobacterium tumefaciens by using incompatibility with a Rhizobium meliloti plasmid to eliminate pAtc58, Plasmid 1985;13:99–105.

[58] Chen S. Ultrafast one-pass FASTQ data preprocessing, quality control, and deduplication using fastp, Imeta 2023;2:e107.

[59] Schmidt T, Friehs K, Flaschel E. Structures of Plasmid DNA. In: Schleef M (editor). Plasmids for Therapy and Vaccination. Hoboken: Wiley-VCH; 2008. pp. 29–43.

[60] Johnson PH, Grossman LI. Electrophoresis of DNA in agarose gels. Optimizing separations of conformational isomers of double- and single-stranded DNAs, Biochemistry 1977;16:4217–4225.

[61] Serwer P, Allen JL. Conformation of double-stranded DNA during agarose gel electrophoresis: fractionation of linear and circular molecules with molecular weights between 3 X 10(6) and 26 X 10(6), Biochemistry 1984;23:922–927.

[62] Slemc L, Jakše J, Filisetti A, Baranasic D, Rodríguez-García A et al. Reference-grade genome and large linear plasmid of *Streptomyces rimosus*: pushing the limits of Nanopore sequencing, Microbiol Spectr 2022;10:e0243421.

[63] Williams LE, Detter C, Barry K, Lapidus A, Summers AO. Facile recovery of individual high-molecular-weight, low-copy-number natural plasmids for genomic sequencing, Appl Environ Microbiol 2006;72:4899–4906.

[64] Germond JE, Vogt VM, Hirt B. Characterization of the single-strand-specific nuclease S1 activity on double-stranded supercoiled polyoma DNA, Eur J Biochem 1974;43:591–600.

[65] Barton BM, Harding GP, Zuccarelli AJ. A general method for detecting and sizing large plasmids, Anal Biochem 1995;226:235–240.

[66] Basta T, Keck A, Klein J, Stolz A. Detection and characterization of conjugative degradative plasmids in xenobiotic-degrading Sphingomonas strains, J Bacteriol 2004;186:3862–3872.

[67] Barbour AG, Garon CF. Linear Plasmids of the Bacterium Borrelia burgdorferi Have Covalently Closed Ends, Science (1979) 1987;237:409–411.

[68] Wibberg D, Blom J, Jaenicke S, Kollin F, Rupp O et al. Complete genome sequencing of *Agrobacterium* sp. H13-3, the former *Rhizobium lupini* H13-3, reveals a tripartite genome consisting of a circular and a linear chromosome and an accessory plasmid but lacking a tumor-inducing Ti-plasmid, Journal of biotechnology 2011;155:50–62.

[69] Goodman MF. The discovery of error-prone DNA polymerase V and its unique regulation by RecA and ATP, J Biol Chem 2014;289:26772–26782.

[70] Remigi P, Capela D, Clerissi C, Tasse L, Torchet R et al. Transient hypermutagenesis accelerates the evolution of legume endosymbionts following horizontal gene transfer, PLoS Biol 2014;12:e1001942.

[71] Christie PJ. Type IV secretion: the Agrobacterium VirB/D4 and related conjugation systems, Biochim Biophys Acta 2004;1694:219–234.

[72] Kubori T, Nagai H. The Type IVB secretion system: an enigmatic chimera, Curr Opin Microbiol 2016;29:22–29.

[73] Lockwood DC, Amin H, Costa TRD, Schroeder GN. The Legionella pneumophila Dot/Icm type IV secretion system and its effectors, Microbiology (N Y) 2022;168.

[74] Nagai H, Kubori T. Type IVB Secretion Systems of Legionella and Other Gram-Negative Bacteria, Front Microbiol 2011;2:136.

[75] Vogel JP, Andrews HL, Wong SK, Isberg RR. Conjugative Transfer by the Virulence System of *Legionella pneumophila*, Science (1979) 1998;279:873–876.

[76] Wagner A, Tittes C, Dehio C. Versatility of the BID Domain: Conserved Function as Type-IV-Secretion-Signal and Secondarily Evolved Effector Functions Within *Bartonella*- Infected Host Cells, Front Microbiol 2019;10.

[77] Costa TRD, Patkowski JB, Macé K, Christie PJ, Waksman G. Structural and functional diversity of type IV secretion systems, Nat Rev Microbiol 2024;22:170–185.

