## Supplementary_figures for "Novel insights into the genome organization of *Rhizobiaceae*: identification of linear plasmids"

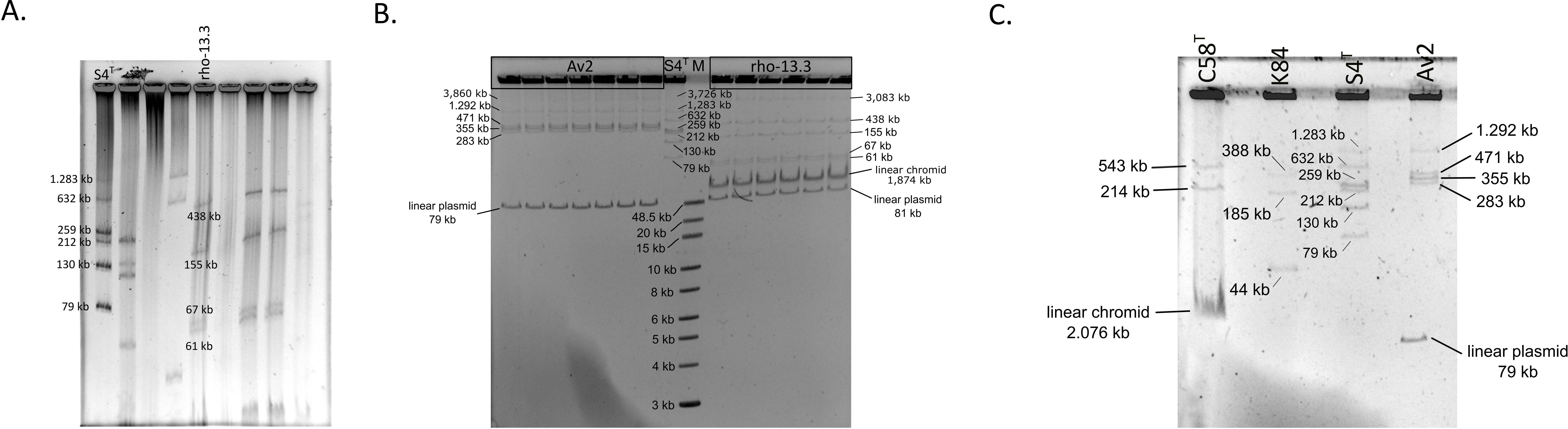


**Fig. S1.** Eckhardt-type agarose gel electrophoresis of *Agrobacterium* sp. rho-13.3 and/or *Allorhizobium* sp. Av2. Gel electrophoresis was performed according the protocol I (**A**) and II (**B** and **C**; see Materials and Methods for more detail). *"Agrobacterium fabrum"* C58$^{T}$, *Allorhizobium ampelinum* S4$^{T}$, and/or *Rhizobium rhizogenes* K84 carrying replicons of known size were used as references. For the gel **B**, a Quick-Load 1 kb Extend DNA Ladder (New England BioLabs, Inc., Ipswich, MA, USA) was included as a marker. A linear chromid of the strain C58$^{T}$ (diffuse band in the gel **C)** is apparently in a sheared state.

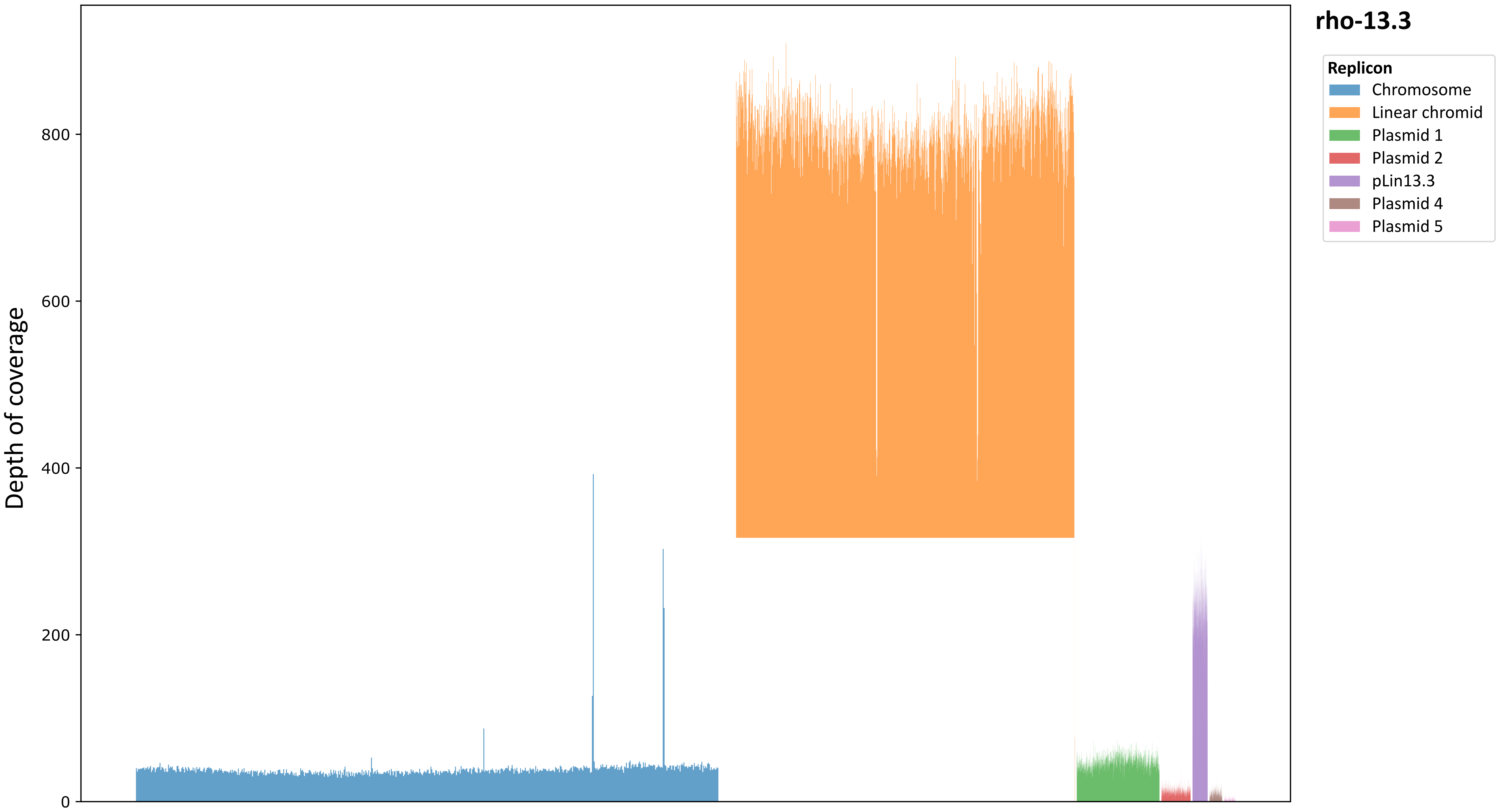


**Fig. S2.** Depth of coverage along replicons. Band corresponding to the linear chromid of strain *Agrobacterium* sp. rho-13.3 was cut from the Eckhardt-type gel, DNA was purified and sequenced using Illumina platform (see Materials and Methods, as well as Table S2 for more detail). Bars correspond to the mean coverage of 5,000 nucleotide windows for the chromosome, 2,000 nucleotide windows for chromid, 500 nucleotide windows for plasmid 1, 200 nucleotide windows for plasmid 2, and 100 nucleotide windows for pLin13.3, and plasmids 4 and 5.

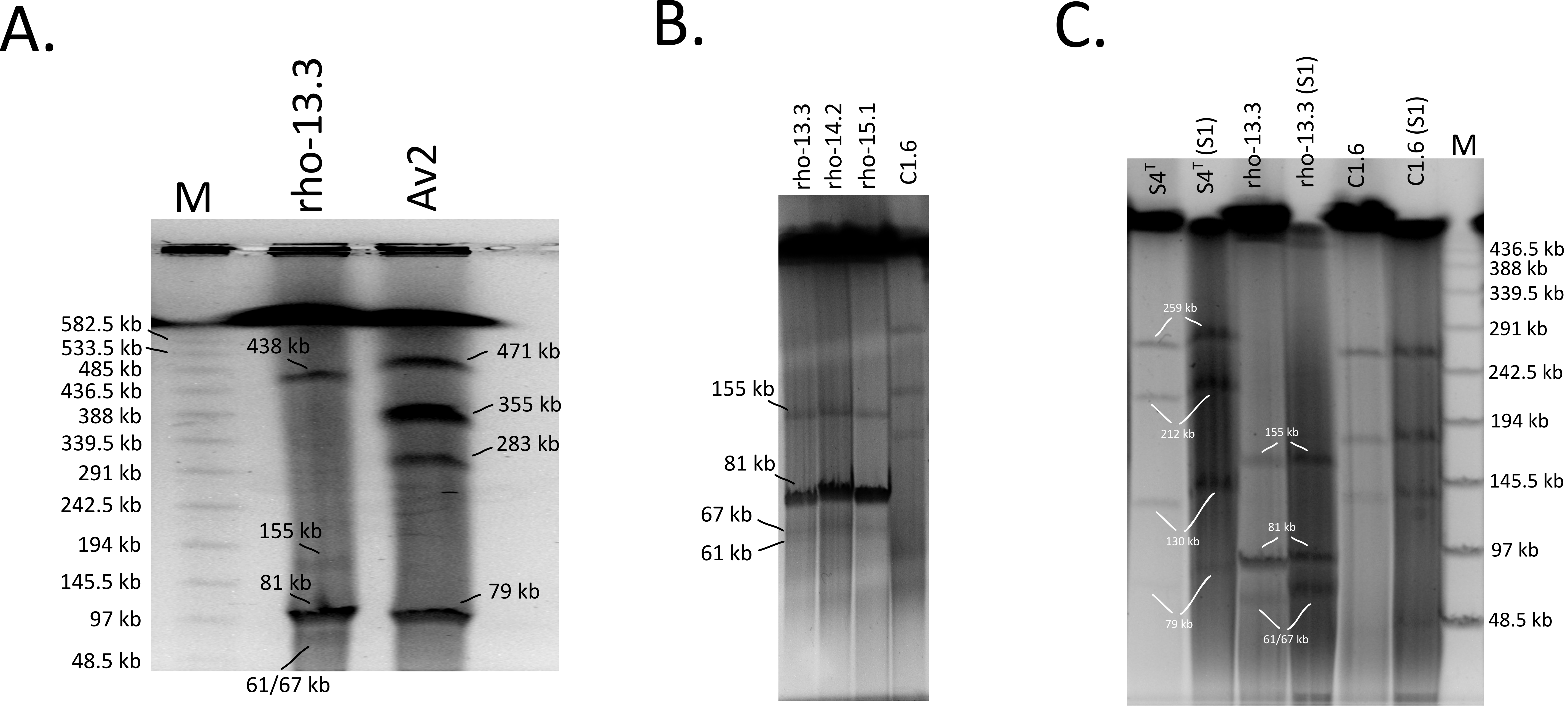


**Fig. S3.** Pulsed field gel electrophoresis (PFGE) of *Agrobacterium* sp. rho-13.3 and/or *Allorhizobium* sp. Av2. Closely related strains rho-14.2 and rho-15.1 were also included in the gel **B**. The gel **C** involved S1 nuclease treatment of agar plugs, as indicated in parentheses. *Allorhizobium ampelinum* $S4^T$, and/or *Agrobacterium* sp. C1.6 were used as references. For gels **A** and **C**, a ProMega-Markers® Lambda Ladders (Promega) was included as a marker. PFGE was carried out in 0.5x TBE buffer using a CHEF-DR III system (Bio-Rad) and the following conditions: voltage 6 V/cm, reorientation angle 120$^o$, switch time 1-25 s (**B**), 5-30 s (**C**) or 10-40 s (**A**), temperature 14°C, time 22 h (**B**) or 30 h (**A** and **C**).

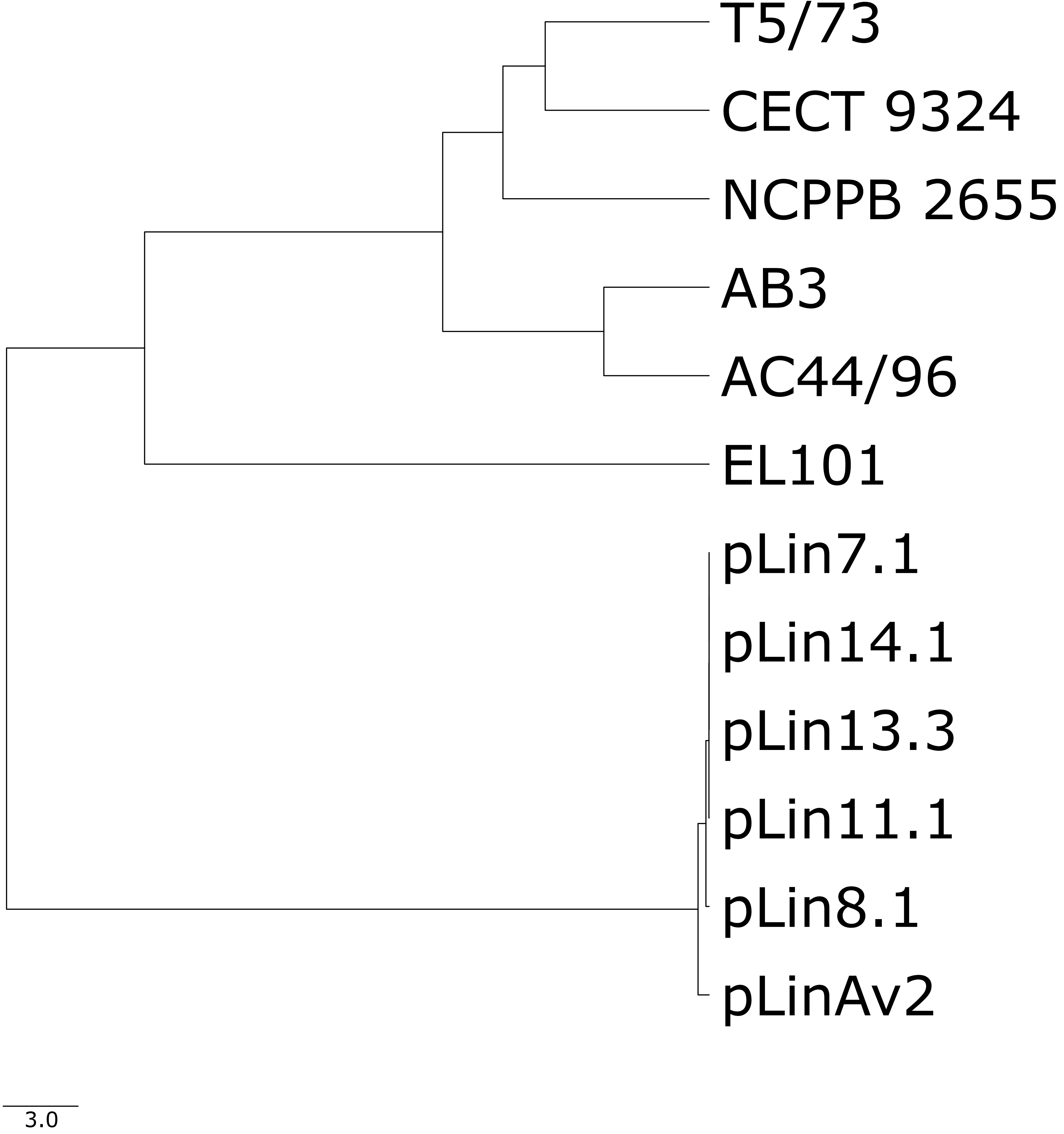


**Fig. S4.** Unweighted Pair Group Method with the Arithmetic mean (UPGMA) hierarchical clustering tree based on pairwise average amino acid identity (AAI) distances between linear plasmids (Tables 3 and S4). AAI values were computed using the EzAAI software (https://github.com/endixk/ezaai). The UPGMA hierarchical clustering tree was constructed with the same software.

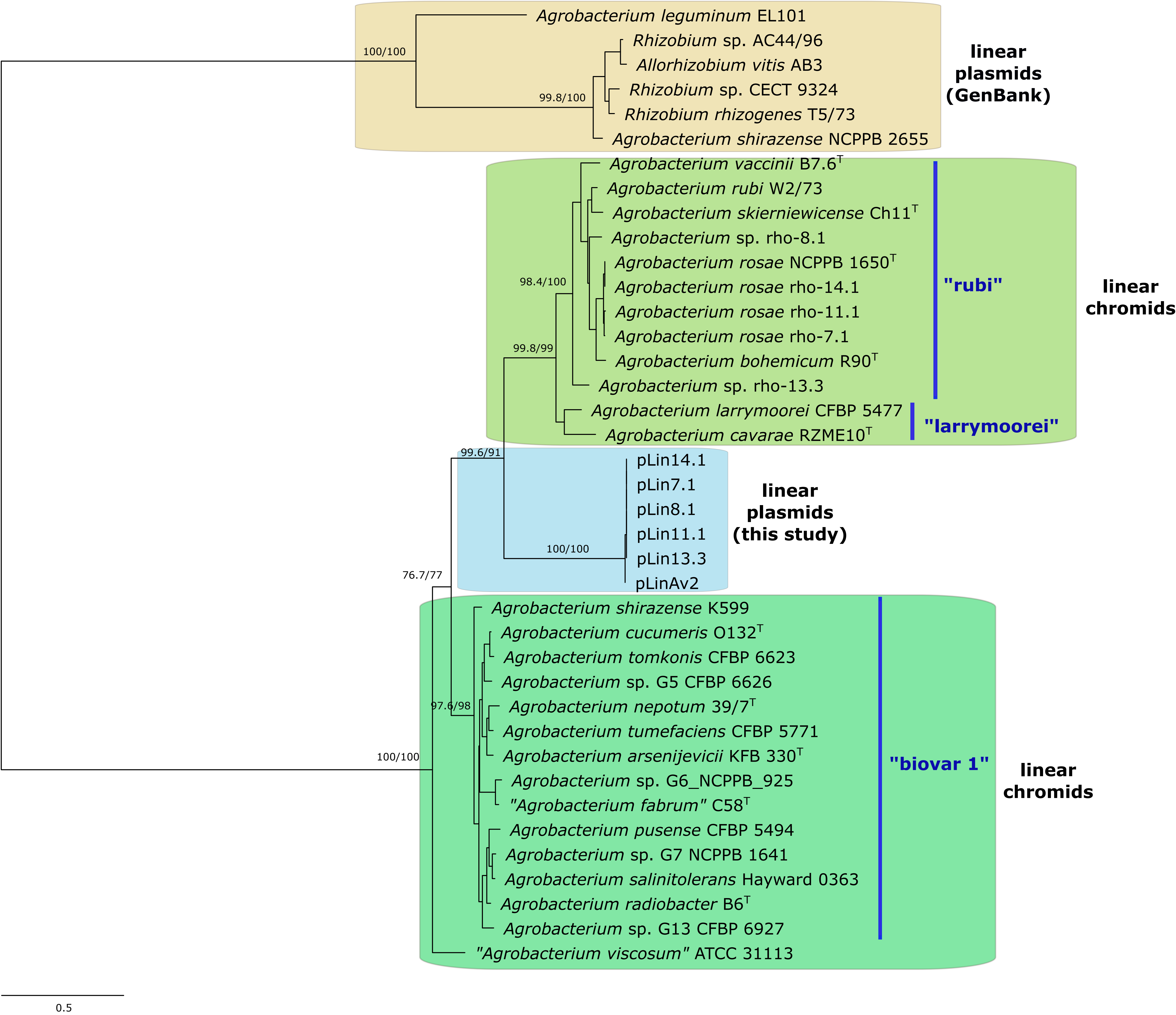


**Fig. S5.** Maximum likelihood tree based on telomere resolvase (protelomerase) protein sequences. *Agrobacterium* sub-clades are indicated in blue-colored text. The tree was constructed using model JTT+G4. The numbers on the nodes indicate the SH-aLRT support values (first value) and ultra-fast bootstrap values (second value). The tree was midpoint rooted. The scale bar represents the estimated number of amino acid substitutions per site.
